# Low extracellular pH protects cancer cells from ammonia toxicity

**DOI:** 10.1101/2024.08.20.608758

**Authors:** Maria Dravecka, Ingvild Mikkola, Terje Johansen, Ole Morten Seternes, Jakob Mejlvang

## Abstract

Ammonia is a natural waste product of cellular metabolism which, through its lysosomotropic ability, can have detrimental effects on various cellular functions. Increased levels of ammonia were recently detected in the interstitial fluid of various tumours, substantiating that high ammonia concentrations are a pathophysiological condition in the tumour microenvironment, alongside hypoxia and acidosis. Since little is known about how cancer cells respond to elevated levels of ammonia in the tumour microenvironment, we investigated how a panel of cancer cell lines derived from solid tumours behaved when exposed to increasing concentrations of ammonia. We found that ammonia represses cell growth, induces genome instability, and inhibits lysosome-mediated proteolysis in a dose-dependent manner. Unexpectedly, we also found that small fluctuations in the pH of the extracellular environment, had a significant impact on the cytotoxic effects of ammonia. In summary, our data suggest that the balance of pH and ammonia within the interstitial fluids of cancerous tumours can have a significant impact on the behaviour and fate of cancer cells.

## Introduction

The development of targeted cancer therapies necessitates the identification of molecular targets which can be based on any specific feature of the cancer that is uncommon in normal cells or tissue. One such feature is represented by the pathophysiological conditions arising in the tumour microenvironment due to the combined effect of cancer-associated metabolic reprogramming and decreased perfusion caused by improper tissue architecture. The best-characterised pathophysiological condition arising in the tumour microenvironment is reduced oxygen availability, known as hypoxia[1]. Hypoxia induces the expression of glucose transporters, glycolytic enzymes, and inhibitory kinases of the pyruvate dehydrogenase complex, which all serve to increase glycolysis in the cancer cells[2,3]. The increased secretion of lactic acid derived from glycolysis, together with increased concentrations of CO_2_ produced from oxidative phosphorylation, causes acidification of the interstitial fluid which is referred to as tumour acidosis[4]. Since both acidosis and hypoxia have a wide impact on the proliferative and metastatic potential of cancer cells[4,5], new therapeutic strategies are being developed to target these two pathophysiological conditions in the tumour microenvironment[6-8].

Besides anaerobic glycolysis, metabolic reprogramming of cancer cells also involves increased dependence on glutamine. Glutamine can serve both as an energy supply and a carbon/nitrogen source for the synthesis of nonessential amino acids, nucleotides and fatty acids[2]. During hypoxia, cancer cells preferentially use glutamine to provide carbon for fatty acid biosynthesis through reductive carboxylation to meet the demand for lipid synthesis[9,10]. The increased flux of glutamine into α-ketoglutarate drives a constant release of ammonia. Although breast cancer cells in vitro have been shown to utilize some of this ammonia as a nitrogen source[11], a positive net production of ammonia is released into the interstitial fluids, causing ammonia concentrations in the tumour microenvironment to reach 1-10 mM[11-13]. Because these concentrations have been shown to reduce viability in various cell lines *in vitro*[14-18], ammonia is emerging as a novel pathophysiological condition of the tumour microenvironment. Yet, there is a paucity of data in the literature addressing how cancer cells are affected by extracellular ammonia.

Ammonia refers to the combined pool of mildly acidic ammonium ions (NH_4_^+^) and weakly basic ammonia gas molecules (NH_3_) which exists in an equilibrium largely governed by pH and temperature. Because NH_4_^+^ is a charged ion, it requires membrane-embedded transporter proteins to cross cellular membranes[19]. In contrast, NH_3_ can freely dissociate over membranes. The different biochemical properties of NH_4_^+^ and NH_3_, cause ammonia to have a lysosomotropic effect, which means it accumulates and alkalises acidic organelles[20]. The cytotoxic effect of ammonia is therefore likely related to its impact on the endolysosomal system which consists of various interconverting acidic vesicles with pH ranging from 6.7 to 4.7, i.e., early and late endosomes, endolysosomes, autolysosomes, phagolysosomes, autophagolysosomes and primary lysosomes[21,22]. For example, the most abundant lysosomal proteases, cathepsins, are only activated when pH reaches a certain lower threshold[23,24]. Consistently, degradation of substrates derived from the endocytic, autophagic, and phagocytic pathways can be reduced when cells are exposed to high levels (≥10 mM) of ammonia[16,20]. In contrast, intermediate levels (1-4 mM) have been suggested to induce autophagy[12,25-27], a process leading to lysosomal degradation of cytoplasm.

Recognising that ammonia levels in the tumour microenvironment may play a significant role in cancer progression, we investigated how different concentrations of ammonia affect various cancer cell lines *in vitro*. We found that ammonia inhibits lysosomal proteolysis in a dose-dependent manner, leading to an accumulation of proteins involved in endocytic, autophagic, and phagocytic pathways. This inhibition results in decreased cell growth, increased cell death and genome instability. Moreover, low extracellular pH, which is often present in the tumour microenvironment, counteracts the effects of ammonia. Therefore, the balance of pH and ammonia within the interstitial fluids of cancerous tumours can therefore have a significant impact on cancer progression and may potentially be targeted in future therapies.

## Results

### Ammonia inhibits cell growth at tumour-relevant concentrations

To investigate how ammonia affected growth of cancer cells, we cultured a panel of human carcinoma cell lines (A549, A431, HT29, and HCT116) in the presence of 0, 4, 10, and 40 mM of NH_4_Cl. To minimize the artificial influx of ammonia from cell metabolism and spontaneous glutamine hydrolysis, we changed the media each day and used media containing GlutaMAX instead of L-glutamine. 4 mM NH_4_Cl moderately repressed growth in all cell lines except HCT116 (Figure 1A). 10 mM NH_4_Cl strongly repressed growth in all cell lines, while cells exposed to 40 mM NH_4_Cl ceased to grow after 24 hours and even caused a small decline in cell numbers over the following two days. To elucidate how ammonia suppresses cell growth, we analysed cell cycle distribution and the extent of DNA synthesis after cells had been cultured for three days in the presence of increasing concentrations of NH_4_Cl. Exposure to 4 mM NH_4_Cl did not have any noticeable effect on cell cycle distribution in any of the cell lines, but slightly reduced the average incorporation of BrdU in A549 cells (Figure 1B-D and S1A). 10 mM NH_4_Cl caused a small increase in cells located in the G1 phase and a decrease in cells undergoing replication (in S and G2/M phase). 10 mM NH_4_Cl also caused an increase in the number of cells in sub-G1 and an increase in cells that contained DNA between 2n and 4n but did not incorporate BrdU (sub-G2). Furthermore, 10 mM NH_4_Cl reduced BrdU incorporation in S-phase cells in all cell lines (Figure S1A). At 40 mM NH_4_Cl, less than 5% of the cells incorporated BrdU and more than 20% of the cells were localized to sub-G1 and sub-G2. To get a more dynamic view of the cell cycle progression we measured how many cells entered M-phase during 6 hours using phosphorylated histone H3 (ser10) as an M-phase marker (Figure S1B). NH_4_Cl exposure caused a dose-dependent decrease in cells reaching M-phase where only very few cells treated with 40 mM NH_4_Cl reached M-phase. We also observed that cell death was induced by NH_4_Cl in a dose-dependent manner leading to substantial cell death (10-40%) in all cell lines at 40 mM NH_4_Cl (Figure 1E). The cell death was not associated with cleavage of PARP or Caspase 3 (Figure 1F and S1C), indicating that NH_4_Cl induces necrosis rather than apoptosis. This conclusion was further supported by an annexin-based apoptosis assay (Figure S1D). We next investigated whether NH_4_Cl exposure activated any DNA damage checkpoints by looking at the phosphorylation status of p53 and H2AX (phosphorylated form referred to as *γ*H2AX). None of the tested concentrations of NH_4_Cl led to increased phosphorylation of p53 at Ser15 in any of the cell lines (Figure 1F and S1C). However, in HT29 and HCT116 cells, 40 mM of NH_4_Cl led to an increase in *γ*H2AX. Lastly, we checked whether ammonia exposure repressed proliferation through the G1/S cell cycle checkpoint mediated by the CDK4/6-Rb signalling hub. Whereas 4 and 10 mM NH_4_Cl did not affect the phosphorylation status of retinoblastoma protein (Rb), hypophosphorylated Rb was detected in cells cultured in the presence of 40 mM NH_4_Cl (Figure 1F and S1C). This hypophosphorylation of Rb coincided with decreased expression of Cyclin D1, substantiating that lethally high concentrations of NH_4_Cl eventually cause cell cycle arrest in G1 through the CDK4/6-Rb signalling pathway. Taken together, this suggests that ammonia inhibits cancer cell growth in a dose-dependent manner, with IC50 values (based on three days of growth) ranging from 2 to 7 mM, and with an onset of cell death with concentrations exceeding 10 mM. Based on the diminished expression of phosphorylated H3 (at serine 10), the lack of ATR/ATM-mediated checkpoint signalling, and the increased prevalence of cells in sub-G2, we hypothesise that ammonia affects DNA replication in a way that causes mitotic catastrophe followed by necrosis. Thus, hyperammonemia in the tumour microenvironment likely contributes to genome instability in residing cancer cells.

**Figure 1.**
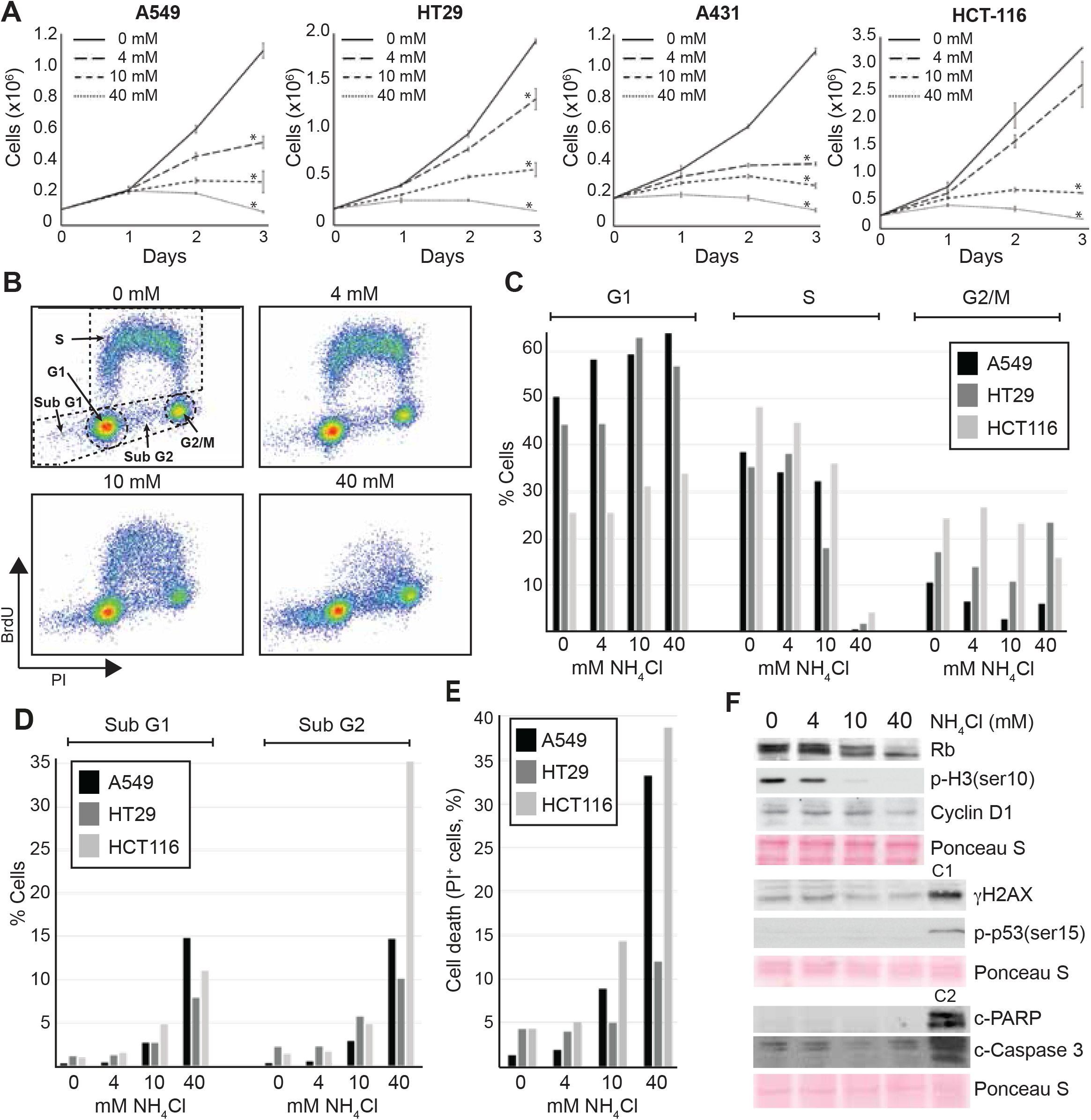
Extracellular NH_4_Cl reduce the growth of cancer cells in a dose-dependent manner. **A**, Growth curves for indicated cell lines cultured in media with indicated concentrations of NH_4_Cl. Error bars display the standard deviation of biological triplicates. *(p < 0.01, two-tailed Students t-test). Source data are provided in the Source Data file. **B**-**D**, Indicated cell lines were cultured in media with indicated concentrations of NH_4_Cl for three days. Then, cells were subjected to 10 minutes of BrdU (20 μM) pulse labelling. Cells were subsequently analysed for BrdU and DNA content (PI, propidium iodide) by flow cytometry. **B**, Dot plot showing DNA content (PI), and BrdU incorporation (BrdU) for HT-29 cells. **C**-**D**, Column chart showing the distribution of cells in each subpopulation defined in B. **E**, Cell death assay. After indicated cell lines had been cultured in media with indicated concentrations of NH_4_Cl for three days, cells were collected by trypsination, stained with propidium iodide, and analysed by flow cytometry (n=20000). **F**, A549 cells were cultured for three days in media with indicated concentrations of NH_4_Cl. Whole-cell lysates were prepared, and Western blot analysis was performed. Ponceau S was used as a loading and blotting control. C1 and C2 represent control lysates. C1; A549 cells cultured overnight in the presence of 50 μM Etoposide. C2; Jurkat cells subjected to heat shock (10 minutes at 44 ° C followed by 6 hours at 37 ° C). Molecular markers and densitometry analysis are shown in the Source Data file.

### Ammonia inhibits lysosome-mediated protein degradation

We observed that NH_4_Cl caused a dose-dependent increase in the number of intracellular granules (Figure 2A) and a dose-dependent increase in cell granularity measured by flow cytometry (Figure 2B), also indicated an increased expression of certain intracellular components. Thus, we decided to perform a mass spectrometry-based quantitative proteomic study to determine how 24-hour exposure to 4 mM NH_4_Cl affected the proteome in A549 cells. Based on three biological replicates, we identified the relative change in expression of 5962 proteins (Figure 2C-D and Datasheet 1A). 208 proteins increased significantly in expression, while 151 proteins decreased (Datasheet 1B-C). Gene ontology enrichment analysis (PANTHER Overrepresentation Test[28]) of all proteins with significantly increased expression, revealed that the cellular components, lysosome, endosome, peroxisome, endoplasmic reticulum, Golgi apparatus, and plasma membrane were significantly enriched (Figure 2E and Datasheet 1D). These data suggest that ammonia predominantly affects the endolysosomal system and that ammonia causes an accumulation of proteins associated with the endosomal system. In agreement, culturing cells in the presence of NH_4_Cl clearly affected lysosomal and endosomal vesicles (Figure S2A). In cells that had not been exposed to NH_4_Cl, the lysosomal marker LAMP1 was localized in small distinct foci, predominantly in the Golgi region, but with increasing concentrations of NH_4_Cl, LAMP1 foci increased in numbers, and their localization expanded to cover the whole cytoplasm. CD63-positive vesicles (endosomes) increased in size with increasing concentrations of NH_4_Cl, and at 40 mM NH_4_Cl, CD63 localized in a very homogeneous population of vesicles with a diameter of approximately 2 μm (Figure S2A). Gene ontology enrichment analysis performed on the list of proteins increasing more than 3-fold identified autophagosomes as one of the most enriched cellular components (Datasheet 1E). In agreement, we found that the canonical marker of autophagosomes, LC3B[29,30], strongly increased in expression with increasing exposure to NH_4_Cl (Figure 2F-G and S2B).

**Figure 2.**
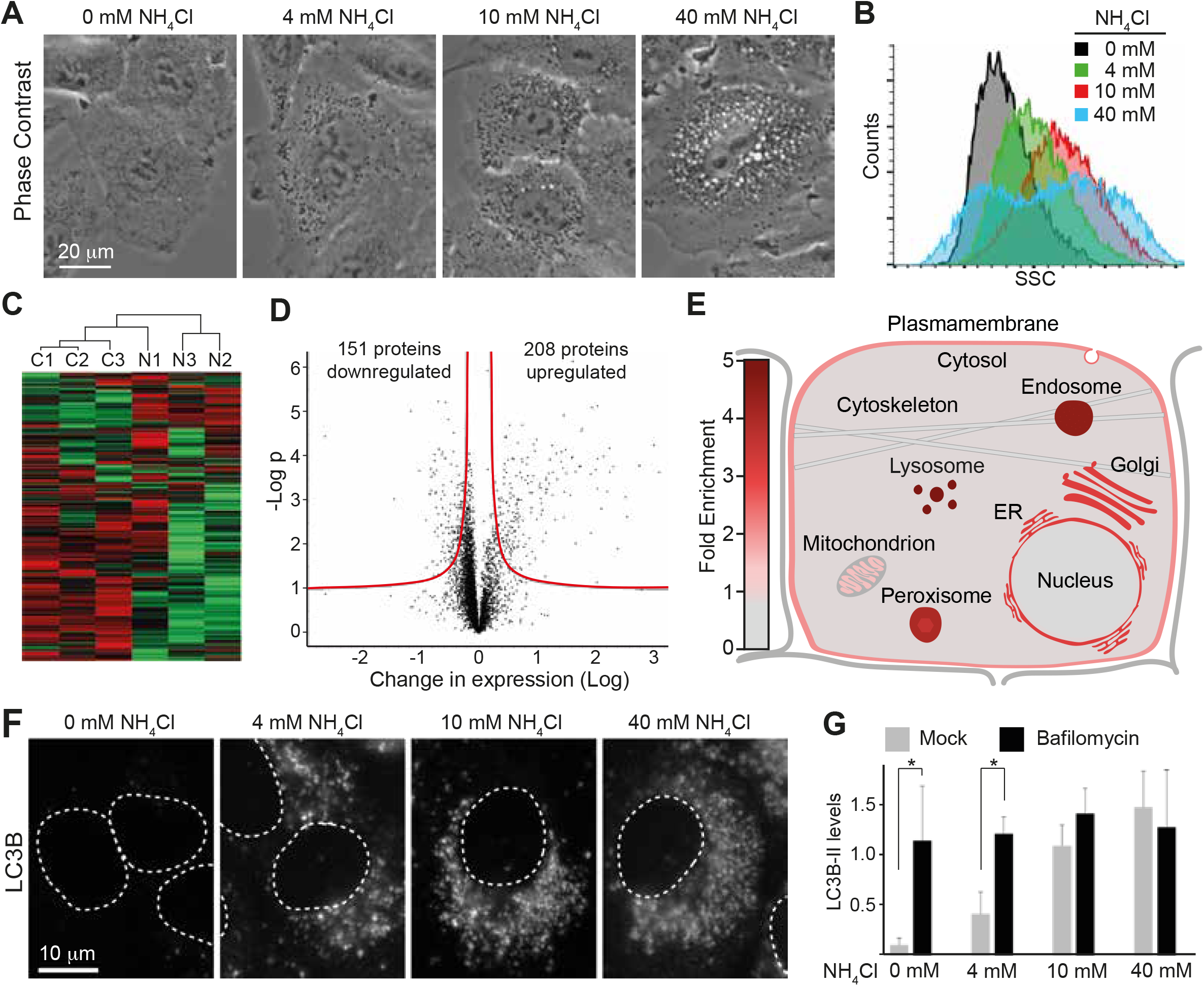
Extracellular NH_4_Cl affects the endolysosomal system. **A**, A549 cells were cultured for two days in media with the indicated concentrations of NH_4_Cl and then imaged in vivo by phase-contrast microscopy. **B**, A549 cells were cultured for three days in the presence of indicated NH_4_Cl. Cells were then analysed by flow cytometry. SSC; side scatter, n=20.000). **C**-**E**, Mass spectrometry-based quantitative proteomics was performed on A549 cells cultured for 24 hours in the absence/presence of 4 mM NH_4_Cl. **C**, Heatmap representation of differentially expressed proteins with hierarchical clustering of samples (control samples: C1-3, 4 mM NH_4_Cl samples: N1-3). Shades of red indicate up-regulated proteins; shades of green indicate down-regulated proteins. **D**, Volcano plot of all identified proteins. The red line depicts the border of significance (see Material and Methods). **E**, Depiction of fold enrichment of displayed cellular components based on gene ontology enrichment analysis. **F**, A549 cells were cultured for two days in media with the indicated concentrations of NH_4_Cl. Cells were then fixed with MeOH, stained by immunohistochemistry, and analysed by fluorescence microscopy. Dotted lines depict nuclei. **G**, A549 cells were cultured for 5 hours in media with the indicated concentrations of NH_4_Cl, either with or without bafilomycin (200 nM). The figure depicts relative LC3B-II levels based on quantitative densitometric analysis of Western blots. Error bars represent the standard error of 5 biological replicates. *(p < 0.05, as determined by two-sided Welch’s t-test). Source data are provided in the Source Data file.

LC3B is conjugated to phosphatidylethanolamine and anchored in the autophagosomal membrane and serves as a recruitment platform for various cargo receptors which add selectivity to the autophagic process[31]. This latter function is not restricted to macroautophagy (formation and processing of autophagosomes) but is also operative in endosomal microautophagy[32]. LC3B, and other ATG8 family proteins, are also conjugated to single membranes in non-canonical autophagy processes that are triggered by a variety of stimuli[33]. Interestingly, the increase in LC3B expression was detected already after 1-2 hours of exposure and continued to increase for at least 24 hours (Figure S2C-D). In contrast to LAMP1 and CD63, LC3B was hardly detectable by immunofluorescence when cells were cultured in media without ammonia (Figure 2F). However, by adding as little as 1 mM NH_4_Cl to the growth media, LC3B puncta appeared (Figure S2E), and with increasing concentrations of NH_4_Cl, LC3B puncta increased in numbers (Figure 2F). These LC3B puncta had a strong tendency to localize to areas positive for CD63, and analysed by high magnification, LC3B puncta seemingly localised within CD63-positive vesicles (Figure S3). The increase in LC3B did seemingly not reflect a stimulation of macroautophagy. Firstly, because the formation of LC3B puncta stimulated by NH_4_Cl was not inhibited by pharmacological (LY294002) inhibition of macroautophagy (Figure S4A), and secondly, because NH_4_Cl did not cause any increase in WIPI2 puncta, an early marker for autophagosome formation[34] (Figure S4B-C). As high extracellular concentrations (≥10 mM) of ammonia have previously been shown to inhibit lysosome-mediated proteolysis[20,29], we next tested how different concentrations of ammonia inhibited the basal lysosome-mediated degradation of LC3B. To this end, we compared how much LC3B increased in expression when cells were exposed to NH_4_Cl, to how much it increased when lysosomal-mediated degradation was completely blocked by Bafilomycin A1. Exposing cells to ammonia alone, led to a dose-dependent increase in LC3B expression (Figure 2G). Concentrations of both 10 and 40 mM NH_4_Cl resulted in an increase comparable to that caused by Bafilomycin A1, indicating that these concentrations blocked lysosome-mediated degradation. This demonstrates that the lysosome-mediated degradation of LC3B gradually decreases with increasing concentrations of NH_4_Cl until it ceases when extracellular NH_4_Cl reaches a threshold (approximately 10 mM). Taken together, our results therefore corroborate that cellular components of the endolysosomal system accumulate in cells exposed to ammonia because ammonia suppresses lysosome-mediated degradation of these components in a dose-dependent manner.

### Low extracellular pH counteracts the lysosomotropic effect of extracellular ammonia

In addition to LC3B, we also observed an accumulation of its associated cargo receptors p62, NBR1, TAXBP1, CALCOCO2 and the receptor for ferritinophagy, NCOA4 (Datasheet 1B). Since we previously identified these proteins to be substrates for starvation-induced endosomal macroautophagy[32], we tested how intermediate concentrations of ammonia affected this process. Cells cultured for 24 hours in the presence of 4 mM NH_4_Cl were starved for amino acids and serum (in the continued presence of 4 mM NH_4_Cl) using either Earle’s balanced salt solution (EBSS) or Hanks’ balanced salt solution (HBSS). Much to our surprise, starvation in EBSS had minimal effect on the expression of LC3B, NBR1, p62, and NCOA4 (hereafter referred to as autophagic substrates), while HBSS caused a substantial reduction (Figure 3A). An inspection of the chemical composition of the two starvation buffers revealed that they mainly differed in the content of sodium bicarbonate (0.35 g/L in HBSS vs. 2.2 g/L in EBSS), which is instrumental for buffering pH. We therefore compared the pH of EBSS and HBSS after they had been acclimatized to 5% CO_2_ and 37°C. This comparison disclosed that while EBSS had a pH of 7.5, HBSS had a much lower pH of 6.8. When we supplemented HBSS with sodium bicarbonate to reach 2.2 g/L, the pH increased to 7.5, and we no longer observed a degradation of the substrates (Figure S4D). Moreover, lowering the pH of the media to 6.8 caused a similar stimulation of lysosome-mediated degradation of autophagic substrates compared to HBSS in cells exposed to 4 mM NH_4_Cl (Figure 3B). Of note, this degradation was minimally affected by pharmacological (LY294002 and SAR405) inhibition of macroautophagy. We next tested whether the accumulation of autophagic substrates upon NH_4_Cl exposure was affected when the pH of the full media was lowered to 6.8. In sharp contrast to cells cultured at pH 7.5, cells cultured at pH 6.8 did not accumulate autophagic substrates when exposed to up to 10 mM NH_4_Cl (Figure 3C and S4E). However, at 40 mM, autophagic substrates accumulated almost to the same level as for cells cultured at pH 7.5. Western blot analysis of one of the central lysosomal proteases, Cathepsin D, revealed that the maturation of Cathepsin D was inhibited by NH_4_Cl in a dose-dependent manner when cells were cultured at pH 7.5. When cells were cultured at pH 6.8, the impact of NH_4_Cl on Cathepsin D maturation was strongly reduced. The proteolytic maturation of Cathepsin D takes place inside endolysosomal vesicles when these reach approximately pH 5[23,24]. The impaired maturation of Cathepsin D could therefore reflect that NH_4_Cl exposure indirectly impairs Cathepsin D maturation by preventing proper acidification of endolysosomal vesicles. To test this hypothesis, we stained living cells cultured at pH 7.5 with the fluorescent dye Acridine Orange. As a monomer, Acridine Orange emits green light (488 nm), but when it is protonated in an acidic environment, Acridine Orange forms aggregates that emit red fluorescence (570 nm)[35]. In the absence of NH_4_Cl, red light was emitted from small puncta in the cytosol, and green light was emitted from nuclei/nucleoli due to its affinity for DNA and RNA (Figure 3D). With the increasing presence of NH_4_Cl, the red puncta in the cytoplasm became both larger and dimmer and started to emit green light (see inserts 1 and 2 in Figure 3D). In contrast, when cells were cultured in media with pH 6.8, concentrations up to 10 mM NH_4_Cl barely had any effect on the Acridine Orange staining pattern. Lastly, we questioned whether the growth inhibitory effects of NH_4_Cl were also lessened when cells were cultured in media with pH 6.8. As previously reported[36], lowering the pH of the growth media caused a small decrease in growth (from a 2.06-fold increase per day (±0.18) to a 1.65-fold increase per day (±0.21)). Whereas 4-10 mM NH_4_Cl suppressed the growth of A549 cells cultured at pH 7.5, exposure to the same concentrations of NH_4_Cl in media with pH 6.8 had no significant growth inhibitory effect (Figure 3E-F). However, when the concentration increased to 40 mM, cell growth was inhibited to the same degree in media with pH 6.8 compared to media with pH 7.5. Similarly, lowering the pH of the growth media significantly improved the growth of A431 and HT-29 cells exposed to up to 10 mM NH_4_Cl (Figure 3G-H).

**Figure 3.**
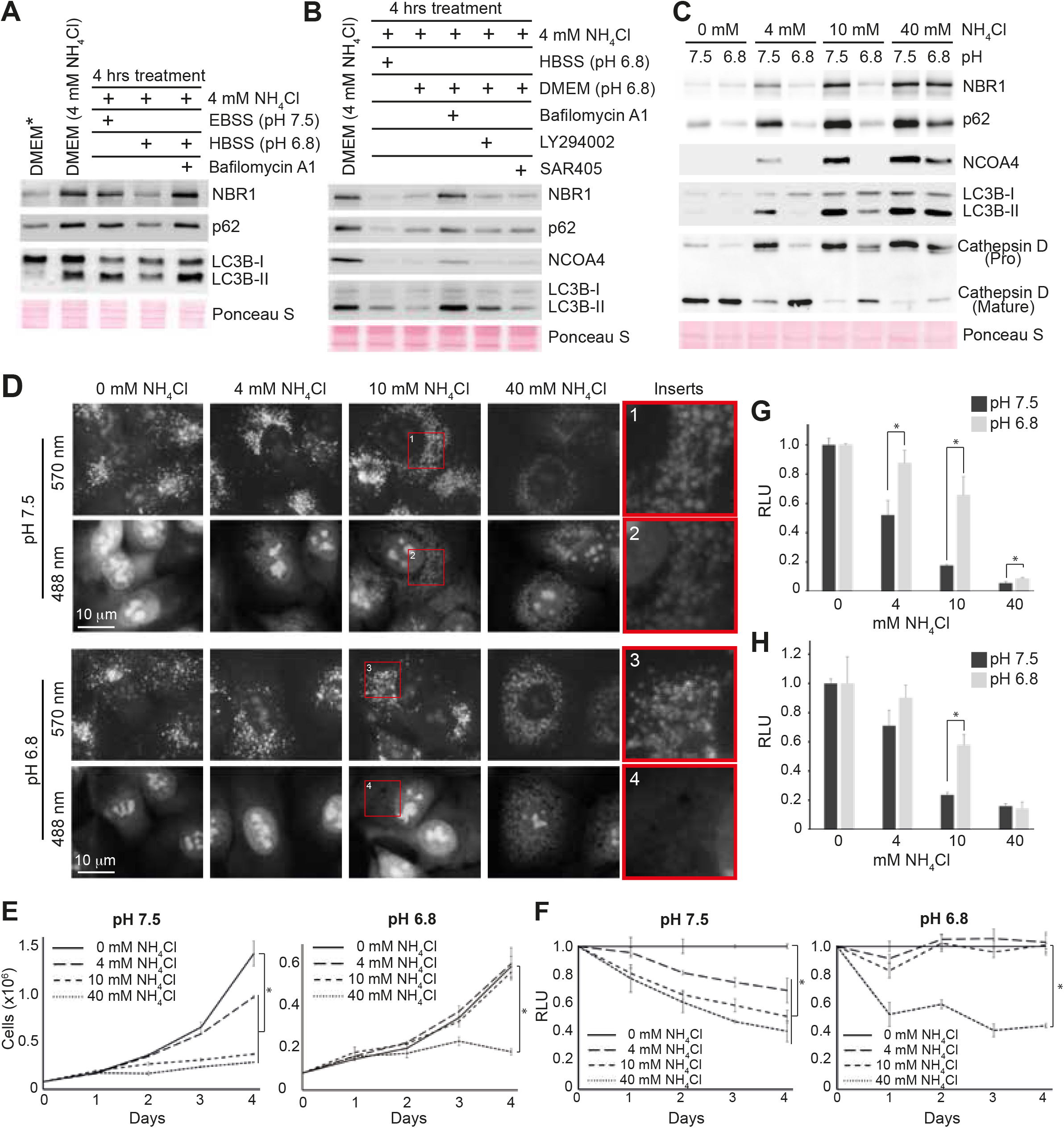
Low extracellular pH reduces the growth inhibitory effect of extracellular NH_4_Cl. **A**-**B**, A549 cells were cultured for one day in media (DMEM) with 4 mM NH_4_Cl and then subjected to indicated treatments for 4 hours. DMEM* represents cells cultured in DMEM without NH_4_Cl. Western blot analysis of indicated proteins. Bafilomycin A1, 200 nM. LY294002, 10 μM. SAR405, 1 μM. Molecular markers and densitometry analysis are shown in the Source Data file. **C**, A549 cells were cultured for two days in media with the indicated pH and indicated concentration of NH_4_Cl. Western blot analysis of indicated proteins. Molecular markers and densitometry analysis are shown in the Source Data file. **D**, A549 cells were cultured for two days in media with the indicated pH and indicated concentration of NH_4_Cl. Then, cells were stained with Acridine Orange for 5 minutes and subsequently analysed by fluorescence microscopy (in vivo). Red squares depict the regions that are magnified to the far right (inserts). **E**, Growth curves based on cell counting. A549 cells were cultured in media with either pH 7.5 or 6.8 and with the indicated concentrations of NH_4_Cl. Error bars represent the standard deviation of 3 replicates. *(p < 0.05, two-tailed Students t-test). **F**, Cell viability assay (CellTiter-Glo). A549 cells were exposed to indicated concentrations of NH_4_Cl while being cultured in media with either pH 7.5 or 6.8. Error bars represent the standard deviation of 3 replicates. *(p < 0.05, two-tailed Students t-test). **G**-**H**, Cell viability assay. HT29 (G) and A431 (H) cells were cultured in media with either pH 7.5 or 6.8 with the indicated concentrations of NH_4_Cl. After three days cell viability was measured by CellTiter-Glo. *(p < 0.05, two-tailed Students t-test). (E-H), Source data are provided in the Source Data file.

In summary our data corroborate that at pH 7.5, extracellular ammonia in the form of NH_3_ diffuses down a pH gradient into the more acidic cytosol and then into the even more acidic intracellular compartments where it becomes protonated (for a more elaborate explanation, see Figure 4). This ion-trapping effect leads to alkalization of the acidic intracellular organelles such as endolysosomal vesicles. In a dose-dependent manner, the alkalization impairs the maturation and function of endolysosomal proteases, thereby affecting the turnover of endolysosomal vesicles, which eventually leads to an accumulation of semi-acidified vesicles. The vesicles increase in size (swell) due to osmotic pressure caused by an accompanying increase in chloride ions[37]. In combination, this affects the many various processes mediated by the endolysosomal system, including energy and amino acid sensing, signal transduction, ionic homeostasis, mitosis, and lysosome exocytosis[38,39], in addition to protein synthesis[20,40], consequently resulting in reduced growth and vitality (Figure 4). Interestingly, our results show that the cytotoxic effect of ammonia is inhibited when the extracellular pH is lowered to pH 6.8. We envision that this effect is due to two specific reasons. Firstly, because a lower extracellular pH will shift the NH_4_^+^/NH_3_ equilibrium in favour of NH_4_^+^, leading to a decreased concentration of extracellular NH_3_. For example, 2.2% of ammonia will be in the form of NH_3_ at pH 7.5, in contrast to 0.4% at pH 6.8 (based on the Henderson–Hasselbalch equation). Secondly, an extracellular pH lower than the intracellular pH results in a reversed pH gradient between the two compartments, thus reducing the ion-trapping effect[8]. The model of how ammonia increases the pH of acidic intracellular compartments is far from novel[15], but the finding that a relatively small decrease in extracellular pH has a tremendous effect on ammonia’s cytotoxicity is new. Consistent with this discovery, the cytotoxic effect of the drug chloroquine, which also increases the pH of endolysosomal vesicles by lysosomotropism, was recently shown to be diminished when the extracellular pH was lowered to 6.8[41,42].

**Figure 4.**
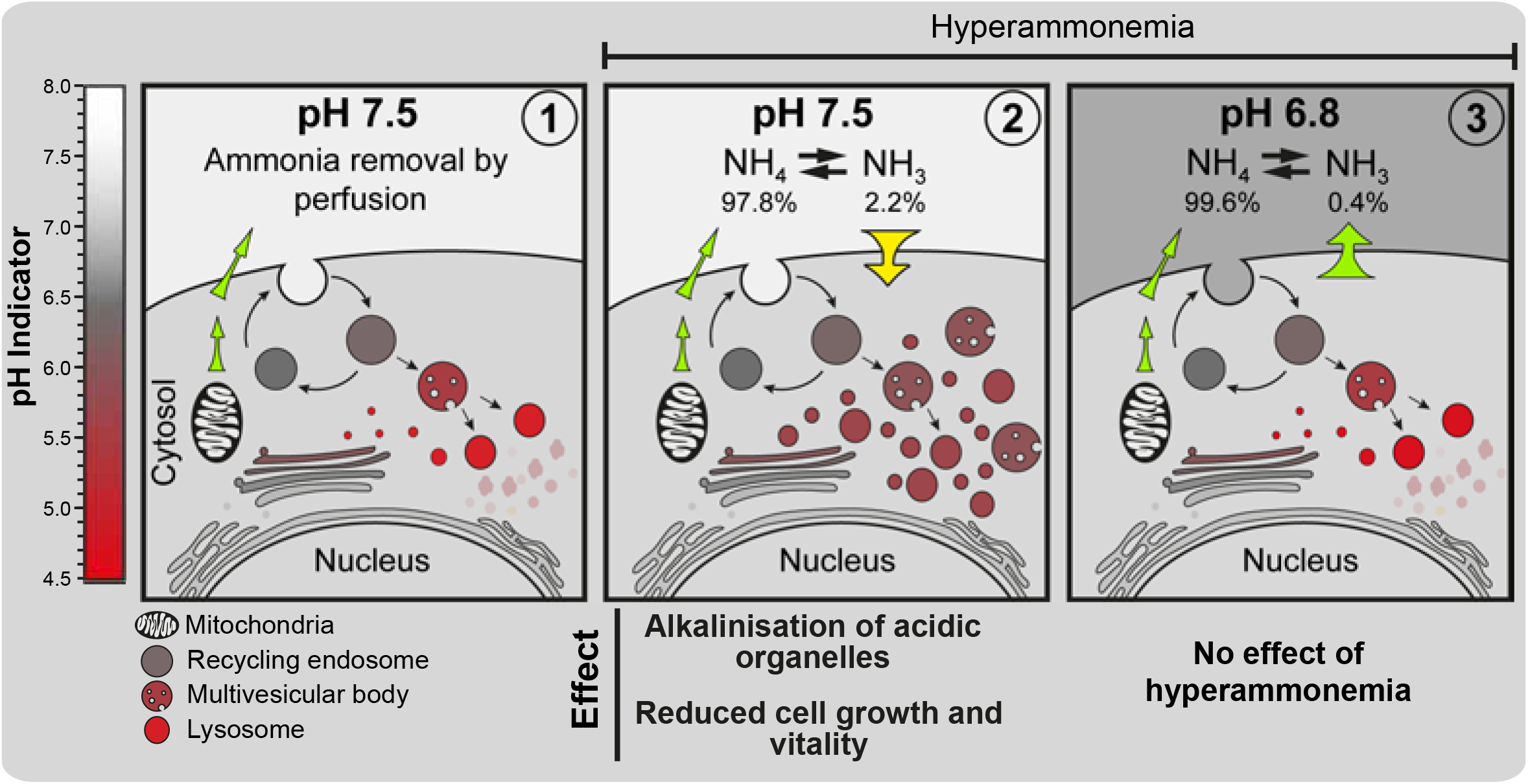
Model summarizing the effect of hyperammonemia at normal physiological pH (pH 7.5) versus low extracellular pH (pH 6.8). **Panel 1**. Increased glutaminolysis leads to a positive net production of NH_4_^+^, which is exported from the mitochondria to the extracellular space (green arrows). When perfusion of the interstitial fluid is normal, ammonia is constantly removed from the microenvironment. **Panel 2**. When perfusion is abrogated, ammonia accumulates in the extracellular space. Under normal physiological pH, 2.2 % of the total ammonia pool will be in the form of NH_3_. As an alkaline gas, NH_3_ can traverse the plasma membrane and enter the more acidic cytosol (yellow arrow). When the influx of NH_3_ exceeds the efflux of NH_4_^+^, ammonia will accumulate in the cytosol, where it leads to a dose-dependent alkalinization of acidic organelles, which impedes their turnover and function. This indirectly represses cell growth and vitality. **Panel 3**. When the extracellular pH is 6.8, less (0.4 %) of the total ammonia pool will be in the form of NH_3_. Therefore, less NH_3_ will be available to enter the cytoplasm. In addition, low extracellular pH instigates a reversed pH gradient (large green arrow), impeding the flux of NH_3_ to the cytosol. Consequently, high extracellular ammonia levels are largely effectless when cells are subjected to low extracellular pH.

## Discussion

Our main finding, that ammonia toxicity is highly dependent on extracellular pH, highlights an important caveat in attempts to elucidate the effects of different pathophysiological conditions in the tumour microenvironment through studies focusing on a single, isolated pathophysiological condition. For example, the effect of hypoxia is widely studied *in vitro* by culturing cells in the diminished presence of O_2_ (1%), but with normal levels of CO_2_ (5%), and no addition of lactate[43]. However, hypoxia in the tumour microenvironment occurs in the presence of hypercapnia (elevated CO_2_) and elevated levels of lactic acidosis[6,44], and several studies have actually shown that the hypoxia-induced cellular response is highly influenced by lactic acidosis[45,46]. Similarly, we find that the cytotoxic effect of ammonia is dependent on the extracellular pH. Since acidosis and hyperammonemia likely are spatially linked, we can only conclude that a much more refined understanding of the spatial balance between ammonia and pH is required to predict to what degree ammonia exerts cytotoxicity in the tumour microenvironment. The current estimates of ammonia concentrations in the interstitial fluid of the tumour are limited to average measurements[11-13]. Naturally, this implies that concentrations could be much higher in the areas of the tumour with the lowest perfusion. Several studies have shown that LC3B accumulates in the more hypoxic/acidic regions of the tumour[41,47,48], furthest away from functional capillaries where cancer cell growth is slowest. Indeed, this observation could reflect an increase in hypoxia-induced autophagy[48], but they could equally well suggest that hyperammonemia and acidosis in these areas reach a balance where ammonia inhibits lysosome efficacy and growth of the residing cells. In the most parsimonious view, our results suggest that pharmacological elevations of pH in the tumour microenvironment would increase the cytotoxic effect of ammonia and thereby kill the cancer cells. However, manipulating the pH of the tumour microenvironment naturally will affect non-transformed components of the tumour microenvironment, such as fibroblasts and immune cells. We found that non-transformed fibroblasts reacted similarly to our panel of cancer cell lines in their response to extracellular ammonia (Bj-1 cells , data not shown), and it was recently shown that extracellular ammonia enhances T cell exhaustion in colorectal cancer[13].

In line with lysosomes being the catabolic end stations of endocytic, autophagic and phagocytic pathways, we found that ammonia in a dose-dependent manner caused an accumulation of component involved in these pathways. For example, the expression of transmembrane receptor amyloid precursor protein (APP), which is endocytosed and subsequently processed in the lysosomes[49], increased 5-fold in cells exposed to 4 mM NH_4_Cl (see Datasheet 1B). Consistently, Komatsu and colleagues also found that ammonia represses the degradation of APP[50]. However, we predominantly focused on elucidating how ammonia affected autophagy and found that the lysosomal-mediated protein degradation of LC3B was blocked by 10-40 mM NH_4_Cl in all tested cell lines. Furthermore, intermediate concentrations (1-4 mM), which only mildly affected cell proliferation, inhibited the lysosomal mediated degradation of LC3B in a dose-dependent manner. Several reports[12,25-27] have suggested that intermediate concentrations of ammonia can induce autophagy. Common to these reports is that they predominantly base their conclusions on autophagic flux measurements estimated by the GFP-LC3 cleavage assay. As described in the “Guidelines for the use and interpretation of assays for monitoring autophagy (4th edition)”[29] ^(see Figure 12 therein)^, the GFP-LC3 cleavage assay can easily be misinterpreted, and in mammalian cells, the assay has actually been reported to detect impairment of lysosomal efficacy caused by increased lysosomal pH[51]. In this light, we find that the experimental results of the aforementioned reports, also reflect that ammonia, in a dose-dependent manner, increases lysosomal pH and thereby represses lysosomal proteolysis. We found no evidence suggesting that ammonia increases the flux of LC3B, nor did we find any evidence suggesting that basal macroautophagy contributed to the accumulation of LC3B in our model system. We therefore suspect that the accumulation of autophagic substrates taking place when cells are exposed to intermediate concentrations of extracellular ammonia, is caused by a dose-dependent inhibition of a basal autophagic process other than macroautophagy. Since the investigated substrates have previously been identified as substrates of endosomal microautophagy[32] we foresee that the basal autophagic activity repressed by ammonia foremostly represents basal endosomal microautophagy.

The maintenance of lysosomal pH depends on a dynamic equilibrium of H^+^ influx and efflux as well as counter-ion movement across the lysosomal membrane[52]. The influx is achieved by the V-ATPase, while the efflux is mediated by leakage through the lysosomal cation channel TMEM175[53]. Counter-ion movement is necessary to dissipate the membrane potential arising from the V-ATPase activity, and is mediated by multiple ion channels and transporters in the lysosomal membrane, which allow the influx and efflux of the counter-ions Cl^-^, K^+^ and Na^+ [52]^. Growing evidence suggests that lysosomal pH is not simply maintained in a strict steady state but can be adjusted to fine-tune lysosomal function. For example, amino acid starvation enhances the assembly of the V-ATPase[54,55], and lysosome-associated membrane protein 1 and 2, regulates the leakiness of TMEM175[56]. Similarly, we envision that intracellular ammonia could be instrumental in regulating lysosome function. This is fully consistent with the recent finding that glutaminolysis, through its metabolic release of intracellular ammonia, represses lysosomal degradation by increasing the lysosomal pH[37,57]. Furthermore, SLC12A9 was recently found to facilitate NH_4_^+^/Cl^-^ co-transport from the lysosome lumen to the cytosol[37]. In SLC12A9 knockout cells, both Cl^-^ and metabolically derived NH_4_^+^ accumulated in enlarged, semi-acidified lysosomal vesicles. This demonstrates that ammonia not only affects the lysosomal pH but also causes an osmotic imbalance in endolysosomal vesicles, which directly affects lysosomal trafficking, fusion and fission events[52].

## Supporting information

SourceDataFile

Datasheet1A

## Acknowledgments

The author acknowledges all members of the Cell Signalling and Targeted Therapy lab and Hallvard Olsvik for insightful discussions. We also thanks Stig Norderval (Department of Gastrointestinal Surgery, University Hospital of North Norway) in his assistance. MS-based proteomic analyses were performed by UiT Proteomics and Metabolomics Core Facility (PRiME), a member of the National Network of Advanced Proteomics Infrastructure, funded by the Norwegian INFRASTRUKTUR-program (project number: 295910). J.M. is supported by a grant from the Erna and Olav Aakres foundation. The authors declare no competing financial interests.

## Author contributions

J.M. wrote the manuscript. J.M. and M.D. performed the experiments. All authors contributed to the experimental planning. M.D, T.J., I.M. and O.M.S contributed to editing the manuscript.

## Materials and Methods

### Cell Culture and Chemical Treatments

A549 cells were cultured in DMEM with 1000 mg/L glucose (SIGMA#D5546) supplemented with serum (9%), penicillin (90 U/ml), streptomycin (10 μg/ml), and GlutaMAX (Gibco, 0.9X). In experiments where pH was adjusted to 7.5, A549 cells were cultured in DMEM (SIGMA#D5030) supplemented with serum (9%), penicillin (90 U/ml), streptomycin (10 μg/ml), GlutaMAX (Gibco, 0.9X), glucose (0.9 mg/ml) and sodium bicarbonate (1.29 g/L). In experiments where pH was adjusted to 6.8, A549 cells were cultured in DMEM (SIGMA#D5030) supplemented with serum (9%), penicillin (90 U/ml), streptomycin (10 μg/ml), GlutaMAX (Gibco, 0.9X), glucose (0.9 mg/ml) and sodium bicarbonate (0.3 g/L). In experiments where pH was adjusted to 7.5, A431 cells were cultured in DMEM (SIGMA#D5030) supplemented with serum (9%), penicillin (90 U/ml), streptomycin (10 μg/ml), GlutaMAX (Gibco, 0.9X), glucose (3.9 mg/ml), sodium pyruvate (0.9 mM), and sodium bicarbonate (1.29 g/L). In experiments where pH was adjusted to 6.8, A549 cells were cultured in DMEM (SIGMA#D5030) supplemented with Serum (9%), penicillin (90 U/ml), streptomycin (10mg/ml), GlutaMAX (Gibco, 0.9X), glucose (3.9 mg/ml), sodium pyruvate (0.9 mM) and sodium bicarbonate (0.3 g/L). In experiments where pH was adjusted to 7.5, HT-29 and HCT-116 cells were cultured in McCoy′s 5A Medium (SIGMA#M4892) supplemented with serum (9%), penicillin (90 U/ml), streptomycin (10 mg/ml) and sodium bicarbonate (1.34 g/L). In experiments where pH was adjusted to 6.8, HT-29 and HCT-116 cells were cultured in McCoy′s 5A Medium (SIGMA#M4892) supplemented with Serum (9%), penicillin (90 U/ml), streptomycin (10 μg/ml), and sodium bicarbonate (0.38 g/L). All cell lines were cultured in HERACELL CO_2_-incubator in the presence of 5% CO_2_, and at 37°C. During experiments, the media was changed each day. Before each media change, the media had been acclimatized in the CO_2_-incubator for a minimum of 1 hour to reach the desired pH and temperature. Similarly, were Hanks’ Balanced Salt Solution (HBSS, SIGMA#H8264) and Earle′s Balanced Salts Solution (EBSS, SIGMA#E2888) were acclimatized in the CO_2_-incubator before use. All cell lines were regularly tested for mycoplasma contamination. All cell lines are derived from the American Type Culture Collection (ATCC). Where indicated, cells were treated with 200 nM bafilomycin A1 (Baf), 10 μM LY294002, 250 nM pp242, 1 μM SAR405, 1 μM wortmannin, 1 μM PIK-III, 100 ng/ml nocodazole, 100 nM LysoTracker^TM^ Red DND-99 (Molecular Probes), and 1 μg/ml acridine orange.

### Detection of Apoptotic and Dead (PI+) cells

For apoptosis detection, we used an APC Annexin V Apoptosis Detection Kit with PI (Biolegend). Cells were harvested by trypsinisation, washed twice with PBS (+1%BSA), and resuspended in 200 μl Annexin V Binding Buffer. Then, 5 μl of APC annexin V and 10 μl of Propidium Iodide (PI) solution were added, followed by an incubation for 15 min in the dark. Subsequently, 400 μl of Annexin V Binding Buffer was added before samples were analysed by flow cytometry on a BD LSRFortessa^TM^ Cell Analyzer. Data were processed using FlowJo^TM^ 10 software.

### Viability with CellTiter-Glo

Cells were seeded into 96-well plates and cultured as indicated. Following treatments, the CellTiter-Glo Luminescent Cell Viability Assay (Promega) was used to measure ATP levels, as instructed by the manufacturer.

### Western Blot Analysis

Samples for Western blotting were harvested in 1X SDS buffer (50 mM Tris, pH 6.8, 2% SDS, and 10% glycerol) and boiled for 5 min. Then protein concentrations were measured using BCA Protein Assay Kit (23227; Pierce) and lysates were calibrated. Bromophenol blue and DTT were added to final concentrations of 0.1% and 100 mM, respectively, before samples were boiled and run on SDS-PAGE gels at 120/160V for 90 min. For blotting, a semidry transfer was used at 100 mA per blot for 1 hour using a buffer containing 14 mM glycine, 48 mM Tris, 0.03% SDS, and 0.15% ethanol. Membranes were stained with Ponceau S before blocking in 5% dry milk in 1X PBS-T for 30 min. Incubation with the primary antibody was performed overnight at 4°C. Membranes were washed six times with 1X PBS-T before incubating with the secondary antibody for 1 hour at room temperature. Membranes were washed six times using 1X PBS-T. Membranes were developed using SuperSignal West Femto Chemiluminescent Substrate (34095; Pierce) and SuperSignal West Pico PLUS Chemiluminescent Substrate (34078; Pierce) on Hyperfilm ECL (18 × 24; GE28-9068-37; GE Healthcare) in a Curix 60 (AGFA) or by chemiluminescence detection using the ImageQuant LAS 3000 (GE Healthcare).

### Antibodies

The following primary antibodies were used in these studies: BrdU (BD Pharmingen, B44), LC3B (Sigma; L7543), NBR1 (Santa Cruz, sc-130380), NCOA4 (Sigma, SAB-1404569), p62/SQSTM1 (BD Biosciences, #610833), PARP (Cell Signalling, #9542), Cathepsin D (Abcam, ab75852), Rb (BD Pharmingen, #14001A), p-p53(Ser15) (Abcam, ab5176), p-H3(Ser10), gH2AX (Cell signalling, #2577, Cyclin D (Santa Cruz, Sc-753), cleaved Caspase 3 (Cell Signalling, #9661), CD63 (DSHB, H5C6), LAMP1 (DSHB, H4A3), and WIPI2 (Abcam, ab105459). The following secondary antibodies were used; HRP-conjugated goat anti-rabbit and anti-mouse (BD Pharmingen #554021 and #554002). Alexa Fluor 568- and 488-conjugated antibodies were purchased from Life Technologies and used at a 1:1000 dilution.

### Cell Cycle Analysis with Propidium Iodide and BrdU

For BrdU/PI labelling, cells were pulse-labelled for 10 min with 10 μM BrdU (Sigma#5002). Following trypsinization, cells were fixed in ice-cold 70% ethanol for 1 hour and then washed once with PBS (+1% serum). Cell pellets were treated with 2 M HCl for 20 min at room temperature before three volumes of 0.1 M sodium borate (pH 8.5) were added for 2 min. Then, cells were washed twice with PBS (+1% serum) and incubated with BrdU antibody (1:20, BD Biosciences cat# 347580) for 1 hour. Following two washes with PBS (+1% serum), cell pellets were incubated for 1 hour with Alexa Fluor 488-conjugated anti-mouse in PBS (+1% serum). After two more washes with PBS (+1% serum), cell pellets were incubated with propidium iodide (50 μg/ml) and RNase (0.25 mg/ml) in PBS for 45 min. All incubations were done in the dark. Samples were finally analysed by flow cytometry on a BD LSRFortessa^TM^ Cell Analyzer, and data were processed using FlowJoTM 10 software.

### Immunocytochemistry and Microscopy

Cells grown on round 12 mm coverslips were fixed for 10 min in cold (-20° C) methanol and rinsed twice with cold PBS. They were then incubated in 2% BSA in PBS-T for 15 min at room temperature to block nonspecific binding. Subsequent incubations with the indicated primary antibodies and fluorescence-conjugated secondary antibodies were performed in PBS-T (containing 1% BSA) at room temperature for 1 hour. Coverslips were rinsed four times for 2 min each with PBS-T following both incubations. Coverslips were subsequently stained with DAPI (0.25 ng/ml) and mounted using VECTASHIELD mounting medium.

For microscopy, we used a DeltaVision Elite microscope (GE) running SoftWoRx 1.0 software utilizing either a 60x or 100x objective. Image channels were acquired sequentially using appropriate filter sets for DAPI, Alexa Fluor 568, and 488. Ordinary phase-contrast micrographs were obtained using an AX10 microscope (Zeiss) equipped with a Retiga 6000 monochrome camera. In all experiments, images shown in individual panels were acquired using identical exposure times or scan settings and adjusted identically for brightness and contrast using Photoshop CS5 (Adobe).

### Proteomic Analysis by Mass Spectrometry

A549 cells were cultured in the absence/presence of 4 mM NH_4_Cl for 24 hours. Then, 2 x 10^6^ cells were lysed in 100 µl of lysis solution (1M Urea, 0.5% SDC, and 100 mM TEAB) and sonicated for 25 cycles (1 minute on, 30 seconds off) with 100% amplitude in a cup horn sonicator with a water cooler (cup horn/watercooler: Qsonica. Sonicator: Fisherbrand FB705 sonicator, Fisher). Aliquots of 70 µg protein were taken out, and proteins were reduced by DTT (5 mM) followed by alkylation with iodoaceamide (15 mM). To remove excess IAA, a DTT solution corresponding to a final concentration of 5 mM was added. Protein digestion was performed with a 1:100 Lys-C (Wako, 125-05061) to protein concentration for 5 hours, followed by a 1:20 trypsin (V511A, Promega) to protein concentration for overnight digestion (16 hours). Digestion with Lys-C and trypsin was carried out under gentle agitation at 37 °C. TMT labeling of peptides was performed according to the manufacturer’s protocol (Thermo Scientific, TMT 6 plex Mass Tag Labeling Kits and Reagents). For TMT labeling, half a 0.8 mg tube of TMT label was used to label a 45 µl sample with 50 µg peptides. A test mix of TMT-labeled samples was prepared by pooling 2 µl of each labeled sample. This was analyzed by mass spectrometry, and the total intensity of all TMT tags was calculated. This allowed us to mix a precise 1:1 ratio of all 6 tags for the final experiment. In the final pooled sample of 50 µg peptides, SDC was removed by precipitation, by adding a final concentration of 2.5% formic acid, incubating for 10 minutes at room temperature, then sample was centrifuging at 13000 rpm for 15 minutes. The supernatant was transferred to a fresh protein low-bind tube and evaporated to dryness in a speedvac. Peptides were reconstituted in 0.5% TFA then concentrated and cleaned up using DPX C18 pipette tips (DPX Technologies, XTR tips 10 mg C18AQ 300Å) followed by evaporation in a speedvac. High pH reverse-phase fractionation[58] was performed on an Ultimate 3000 offline HPLC. C18-purified peptides were reconstituted in 20 mM ammonium formate pH 10 and loaded onto an RP column (Waters Acquity UPLC® BEH C18 1.7 µm 2.1 x 100 mm column). The samples were fractionated using a linear gradient of 0-60 % B (90% ACN, 20 mM ammonium formate, pH 10) at 150 µl/min for 30 min. Twenty fractions were collected and pooled into 10 fractions using the mixing strategy Fr1 + Fr11, Fr2 + Fr12, and so on. Fractions were frozen at – 80°C. Samples were then dried in a speedvac and reconstituted in 2 µl 2% ACN 0.1% formic acid.

Peptides were analyzed using a Thermo Scientific Orbitrap Fusion Lumos mass spectrometer coupled to a Thermo Scientific EASY-nLC 1200 nanoLC. Peptides were loaded onto a PepMap C18 EASY-spray column (2 µm, 100Å, 75 mm x 50 cm) (Thermo Scientific). Peptides were fractionated using a 4-72% acetonitrile gradient in 0.1 % formic acid over 140 minutes at a flow rate of 300 nl/min; 4-6% in 5 minutes, 6-32 % in 120 minutes, 32-72 % in 5 minutes, and isocratic for 5 minutes. Eluted TMT peptides were analyzed using an SPS-MS3 method. MS1 was acquired in the Orbitrap (120k resolution with a scan range of 400-1400 m/z, AGC target 4.0 x 10^5^). MS2 was collected in the Ion Trap after CID fragmentation. MS3 was collected in the Orbitrap (HCD, 7.5 K resolution with AGC target 1.5 x 10^5^, and max IT 150). Raw data was analyzed using MaxQuant (version 2.4.9.0)[59] with the integrated Andromeda search engine. MS/MS data was searched against the Uniprot Human database. An FDR of 0.01 was needed to give a protein identification.

Perseus (version 2.0.11) was used for statistical analysis. TMT intensities were log2 transformed. Proteins with less than 3 valid values were filtered out. Missing values were imputed from normal distribution with a down shift of 1.8. Volcano plot was created using a t-test and FDR of 0.05 and S0 as 0.1.

---

**Figure S1.**
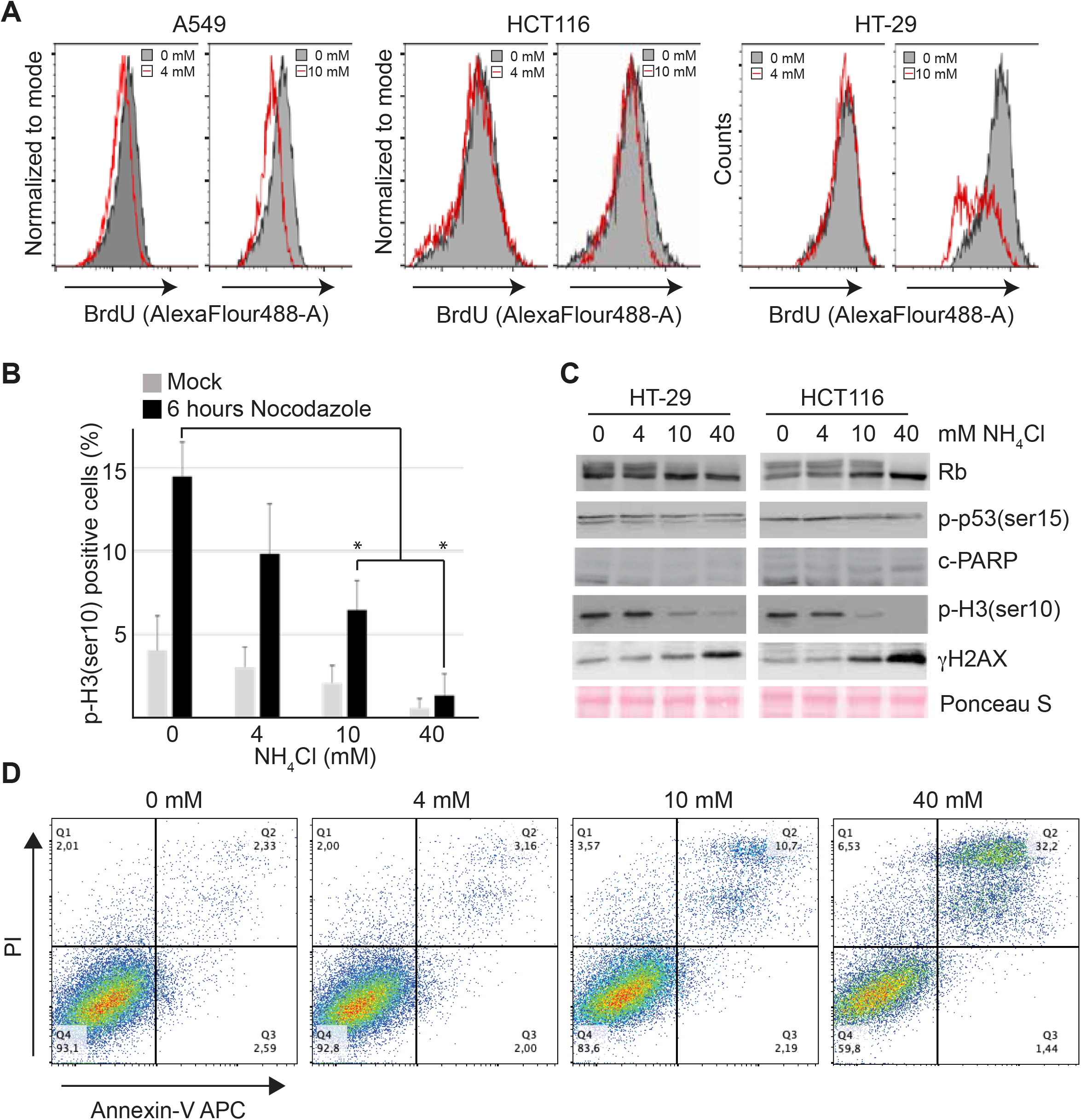
**A**, Indicated cell lines were cultured in media with indicated concentrations of NH_4_Cl for three days. Then, cells were subjected to 10 minutes of BrdU (20 μM) pulse labelling. Cells were subsequently analysed for BrdU and DNA content (PI) by flow cytometry. The histogram shows the level of BrdU incorporation. **B**, A549 cells were grown in the presence of the depicted concentrations of NH_4_Cl for three days. Then, cells positive for Histone 3 phosphorylated at ser10 (p-H3(ser10)) were registered by immunofluorescence. Where indicated, cells were treated with 100 ng/ml Nocodazole to arrest cells in M-phase. Error bars represent the standard error of the mean. *(p < 0.05, as determined by two-tailed students t-test). Source data are provided in the Source Data file.**C**, Western blot analysis of HCT116 and HT29 cells grown for three days in the presence of the indicated concentrations of NH_4_Cl. Molecular markers and densitometry analysis are shown in the Source Data file. **D**, A549 cells were grown in the presence of the depicted concentrations of NH_4_Cl for three days. Then, cells were analysed for apoptosis by Annex-in-V and propidium iodide (PI) staining.

**Figure S2.**
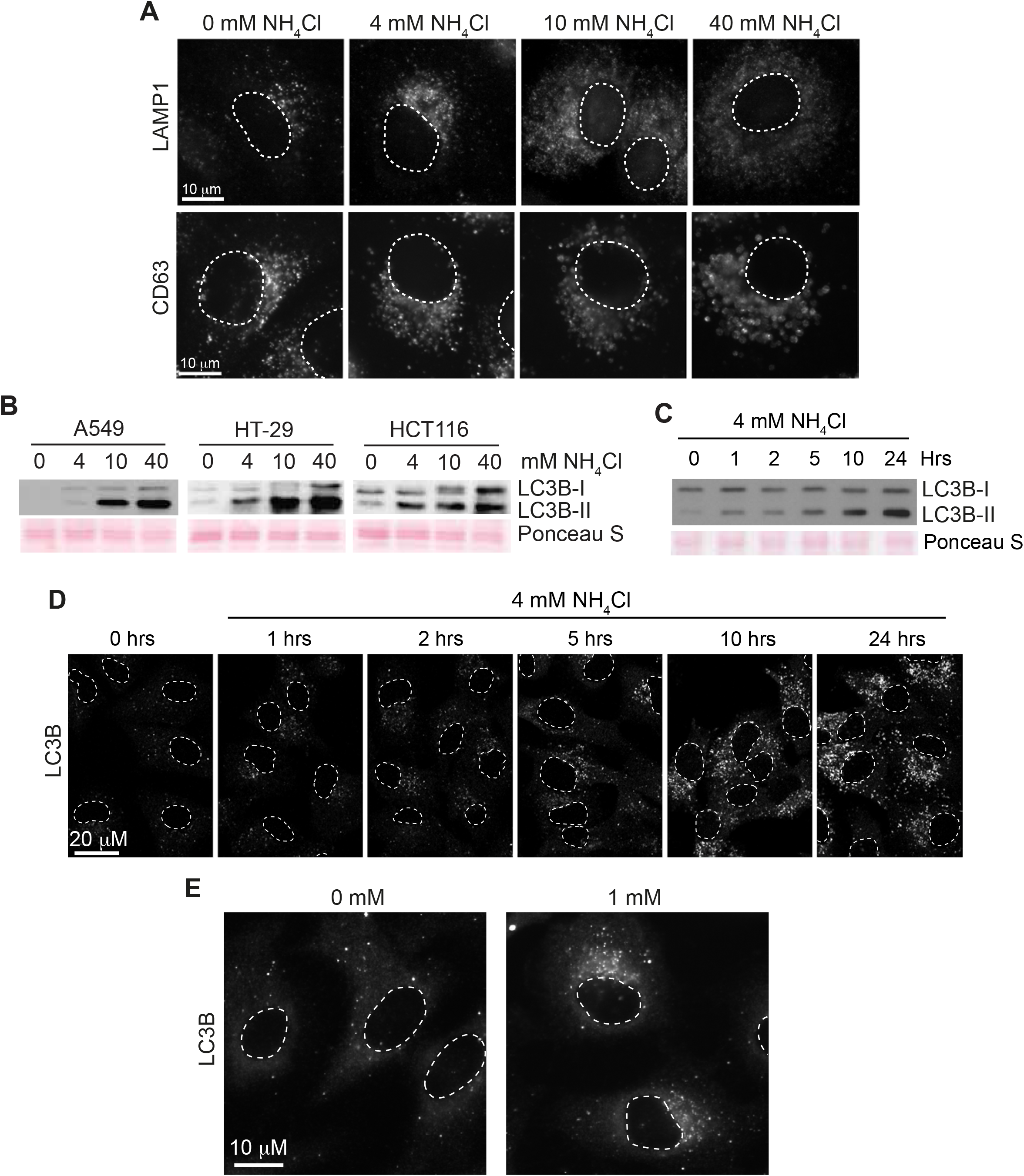
**A**, A549 cells were cultured for two days in media with the indicated concentrations of NH_4_Cl. Then, they were fixed with MeOH, stained by immunohistochemistry, and analysed by fluorescence microscopy. Dotted lines depict nuclei. **B**, Indicated cell lines were grown for three days in the indicated presence of NH_4_Cl at the indicated concentrations. Whole-cell lysates were prepared, and Western blot analysis was performed. Ponceau S was used as a loading and blotting control. Molecular markers and densitometry analysis are shown in the Source Data file. **C-D**, A549 cells were cultured for 0-24 hours in media with 4 mM NH_4_Cl. (**C**), Whole-cell lysates were prepared, and Western blot analysis was performed. Ponceau S was used as a loading and blotting control. Molecular markers and densitometry analysis are shown in the Source Data file. (D), Cells were fixed with MeOH, stained by immunohistochemistry, and analysed by fluorescence microscopy. Dotted lines depict nuclei. **E**, A549 cells were cultured for 24 hours in media with or without 1 mM NH_4_Cl. Then, they were fixed with MeOH, stained by immunohistochemistry, and analysed by fluorescence microscopy. Dotted lines depict nuclei.

**Figure S3.**
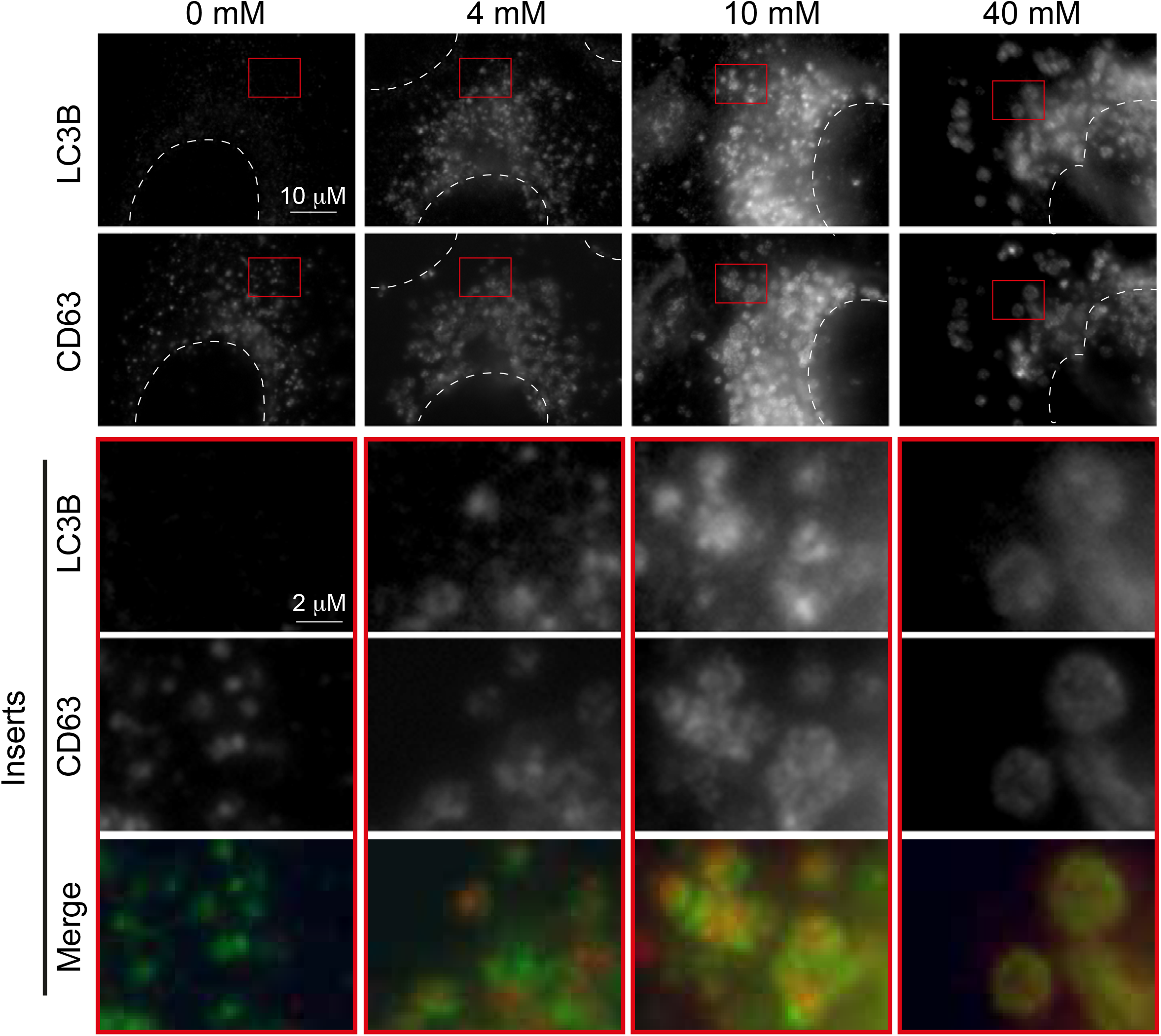
A549 cells were cultured for 24 hours in media with the indicated concentrations of NH_4_Cl. Then, they were fixed with MeOH, stained by immunohistochemistry, and analysed by fluorescence microscopy. Dotted lines depict nuclei. In the merged inserts, green represents CD63 while red represents LC3B.

**Figure S4.**
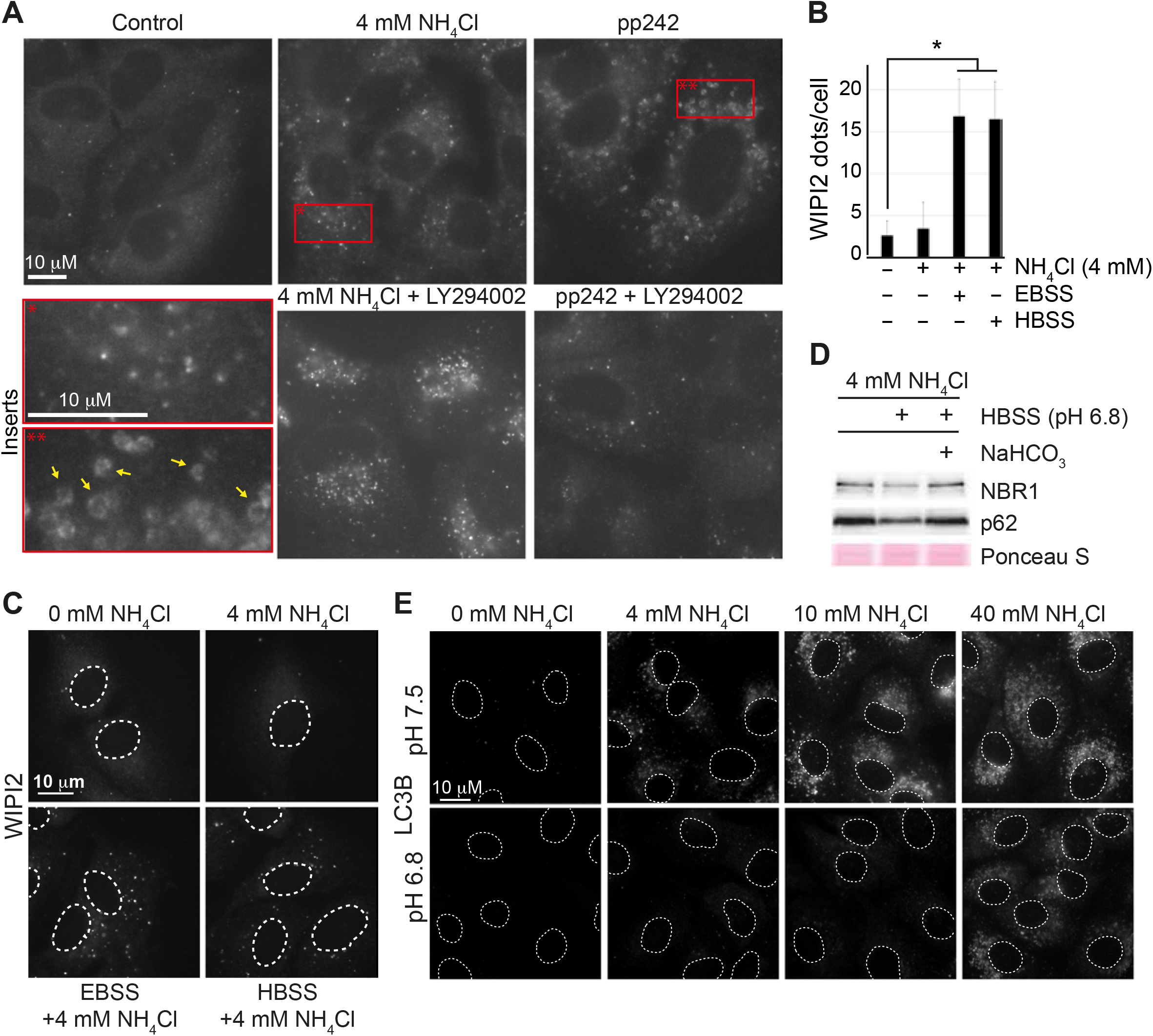
**A**, A549 cells were cultured for 4 hours in the indicated presence of 4 mM NH_4_Cl, 250 nM pp242, and 25 μM LY294002. Then, the cells were fixed with MeOH and stained with an antibody targeting LC3B. Red squares represent the area shown with increased magnification in the inserts. Yellow arrows highlight classical autophagosomes. **B-C**, A549 cells were cultured for one day in media with 4 mM NH_4_Cl. Where indicated, cells were subjected to 4 hours of starvation (HBSS or EBSS) in the continued presence of NH_4_Cl. Then, the cells were fixed with MeOH and stained with an antibody targeting WIPI2. **B**, Quantification of WIPI2 dots per cell. Error bars represent the standard deviation (n>20). *(p < 0.01, two-tailed Students t-test). Source data are provided in the Source Data file. **C**, Representative images of WIPI2 dots. White dotted lines depict nuclei. **D**, A549 cells were cultured for one day in the presence of 4 mM NH_4_Cl and then subjected to the indicated treatments for 4 hours in the continued presence of 4 mM NH_4_Cl. Then, whole-cell lysates were prepared, and Western blot analysis was performed. Ponceau S was used as a loading and blotting control. Where indicated, HBSS was supplemented with NaHCO_3_ to reach a final concentration of 2.2 g/L. This supplementation caused the pH to increase to 7.5. Molecular markers and densitometry analysis are shown in the Source Data file. **E**, A549 cells were cultured for two days in media with the indicated pH and indicated concentration of NH_4_Cl. Then, the cells were fixed with MeOH and stained with an antibody targeting LC3B. Dotted lines depict cell nuclei.

